# RAD: A Read-structure Agnostic Demultiplexer for Single-Cell Long-Read Sequencing and Analysis

**DOI:** 10.64898/2026.05.29.728810

**Authors:** Chinmay M. Vaidya, Margaret C. Carpenter, Leena Abdullah, Fred W. Kolling, Yina H. Huang, Li Song, Margaret E. Ackerman

## Abstract

Single-cell long-read sequencing (LRS) techniques enable the analysis of full transcript sequences within a cell. However, LRS introduces computational challenges for parsing information, often necessitating the development of custom workflows to handle complex read layouts, such as split combinatorial barcodes. To address these challenges, we introduce an error-robust, read-structure agnostic demultiplexer (RAD). In RAD, users can easily specify read structure, such as adapter sequence and relative barcode position, and can rapidly extract these elements for each read. RAD implements highly accurate barcode whitelist generation from long-read data alone, without the benefit of short-read support or priors while uniquely avoiding common but previously uncharacterized errors during demultiplexing. RAD’s comprehensive barcode error correction protocols allow for more sensitive and precise barcode-read assignments. In synthetic and real-world benchmarks against gold-standard demultiplexers, RAD is faster and achieves better biological discovery capability in downstream gene expression analysis than existing pipelines. We show RAD can be applied to datasets ranging from high-definition long-read spatial transcriptomics to targeted long-read single-cell immune receptor sequencing, making it a powerful and scalable tool for both standard and custom long-read sequencing experiments.

## Main

Long-read sequencing (LRS) provides genomic context that traditional short-read sequencing cannot^1–3^. It allows users to differentiate between distinct isoforms and splice variants^4,5^, provides resolution across long repetitive introns^6,7^, and captures full-length transcripts of large genes that would otherwise be truncated^8,9^. The application of LRS to short-read techniques such as targeted single-cell isoform sequencing, combinatorial barcoding, and spatial transcriptomic sequencing offers insight into how transcript diversity may influence cellular identity. LRS experimental designs have expanded in scope and complexity dramatically as the field has evolved, utilizing different and complex arrangements of adapters, barcodes, unique molecular identifiers (UMIs), and additional linkers. However, the biological insights that can be uniquely derived from LRS have been bottlenecked, in part, by challenges in the reliable identification and assignment of reads to their barcode of origin.

Computational processing steps for long-read sequencing can be broadly categorized as either read pre-processing, such as demultiplexing that identifies information like cell barcodes, or read alignment and subsequent annotation. While multiple groups have developed powerful tools and benchmarking datasets for the latter^10,11^, demultiplexing remains both an area of active computational development and a major limitation in the implementation of new LRS techniques and applications. Demultiplexing itself comprises two distinct computational stages: whitelist generation, in which the set of valid cell barcodes is discovered, and per-read barcode assignment, in which each read is corrected and set to a member of that whitelist. In both steps, LRS demultiplexing tools must handle complex error types, such as chimeric and truncated reads, that cannot be discerned or resolved by per-base and read quality metrics^12^. These complex errors may compound during demultiplexing, resulting in both misidentification of barcodes and inadvertent overinflation of unique molecular identifier (UMI) counts due to error-generated singletons^13^. As a result, processing long-read data with existing tools often necessitates or substantially benefits from manual user intervention, paired short-read data, or custom-built demultiplexers for the task at hand. Together, these shortcomings add to the cost, complexity, and time required to analyze and interpret single-cell long-read data. Additionally, as custom LRS pipelines are developed to address novel biological questions^8,14,15^, the complexity of read structures and the computational burden of demultiplexing are likely to increase.

To meet this existing and growing need, we developed a Read-structure Agnostic Demultiplexer (RAD). RAD is a single unified pipeline that processes various complex read structures and multiple sequencing modalities, performing whitelist generation, error correction, and demultiplexing from long reads alone without requiring paired short-read sequencing data. RAD is also long-read platform agnostic, supporting both Oxford Nanopore (ONT) and Pacific Biosciences (PB) sequencing. Protocols with custom read structures require only editing a simple layout CSV that specifies static and variable elements from 5’ to 3’, rather than significant effort in pipeline re-engineering. RAD thus scales with the diversifying landscape of LRS protocols and accepts fully custom layouts alongside bundled templates. This design enables single-pass whitelist generation and demultiplexing of arbitrarily complex read structures, including LR-Split-seq, which presents multiple barcodes interspersed with linkers^8,16^, ONT bulk rapid barcoding, in which barcodes flank both ends of the insert, and Visium 3’ HD, comprising combinatorial adjacent barcodes, without multiple tool invocations or external post-processing. To facilitate downstream alignment and evaluation pipelines, RAD can generate and return custom-formatted read headers after demultiplexing, preserving broad compatibility with pre-existing downstream processing tools.

In this work, we evaluate RAD against the two gold-standard demultiplexers, BLAZE^17^ and Flexiplex^15^, across multiple datasets spanning diverse sequencing chemistries and read structures, benchmarking speed, memory use, and barcode/UMI assignment accuracy across all three tools. We further quantify demultiplexing precision, sensitivity, and propensity to overcorrect in the context of paired short- and long-read single-cell and single-nuclei sequencing. Finally, we highlight the contribution of RAD’s robust demultiplexing in more accurate and confident biological exploration of differential gene expression analysis from long-read single nuclei sequencing. Together, these tools and analyses support broad application of long read sequencing technology to provide new biological insight. RAD is provided as an open-source C++ package at https://github.com/indianewok/rad.

## Results

### Method Overview

We developed a robust, error-tolerant, read-structure agnostic demultiplexer (RAD) for long-read single-cell sequencing data (**Figure 1A**). RAD supports flexible read-layout specification: users define the relative positions of static elements, such as adapters, primers, linkers, and poly-T tails with known sequences, and those of variable elements, such as barcodes, UMIs, and captured reads. It can generate whitelists from LRS data alone without the need for paired short-read sequencing. As part of whitelist generation, RAD utilizes a saddle-point algorithm to separate low-count barcodes from ambient noise and heuristically modifies the resulting threshold depending on other data characteristics. Because of this method of whitelist generation, RAD can rescue error-prone reads, avoid overcorrection, and process data without dependence on paired short-read data. For demultiplexing, RAD first identifies static elements and then uses them as anchors to extract variable elements, parsing each read in both forward and reverse orientations. RAD ships with several pre-configured layout files and complete chemistry whitelists for 10x 3′ and 5′ Chromium, Visium, Visium HD, LR-Split-seq, and bulk Nanopore rapid-barcode chemistries (**Table 1**), but its implementation is flexible to user formats and allows addition of new read structures by manually editing the layout file. During demultiplexing, RAD’s barcode correction continuously operates against the barcode whitelist, accepting a barcode only when independently observed in raw reads. RAD also resolves chimeric concatenated reads, common long-read artifacts in which adapters from multiple distinct molecules ligate end-to-end. Reads with aligned elements in multiple orientations are flagged and subsequently parsed as distinct demultiplexed outputs, thus reclaiming concatenated data that other tools may discard. Outputs can be returned split per valid barcode or pooled in bulk, with user-customized or default headers that are Sequence Alignment Map (SAM) format-compatible.

**Figure 1.**
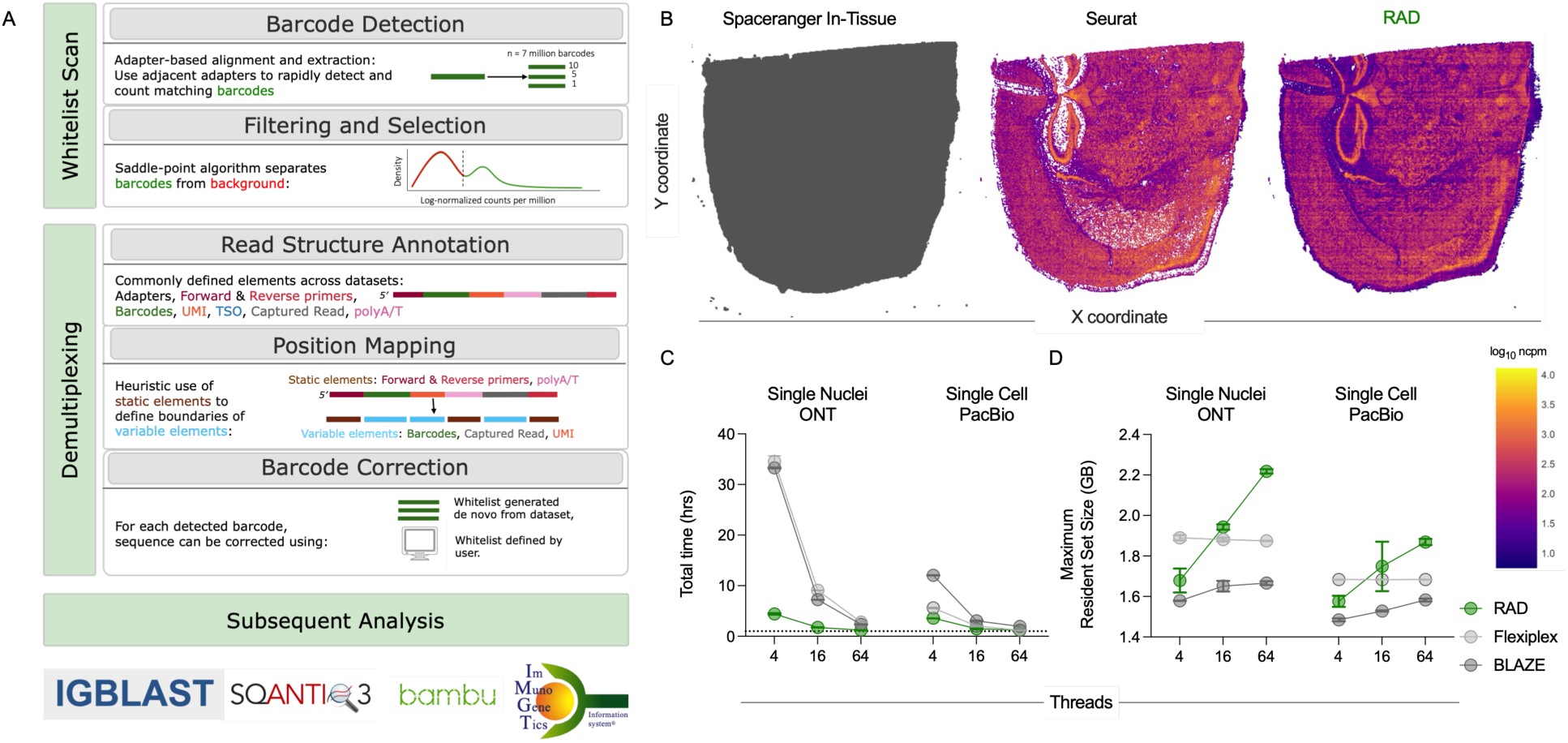
RAD is a flexible, efficient, and broadly applicable tool for demultiplexing single-cell long read sequencing data. **A**. A schematic of RAD workflow. **B**. Visualization of valid barcode spots (gray) determined by SpaceRanger (*in_tissue* metadata of gene expression output) from Visium HD spatial transcriptomic data (left) and log-transformed normalized read counts (log_10_ ncpm) per 16-mm spot for Seurat (center) and RAD (right) processed data. **C-D**. Total processing time (**C**) and maximum memory usage (**D**) for single-cell Oxford Nanopore Technologies (ONT) and single-cell Pacific Biosciences (PB) LongBench datasets.

**Table 1.** Built-in read layout templates in RAD.

| Layout key | Assay / Modality | Barcode element(s) | Barcode length (nt) | UMI length (nt) | Default whitelist kit(s) | Whitelist size |
|---|---|---|---|---|---|---|
| three_prime | 10x Genomics 3' scRNA-seq | barcode | 16 | 10 | 10x_3v3 | 6,794,880 |
| five_prime | 10x Genomics 5' scRNA-seq | barcode | 16 | 10 | 10x_5v1 | 737,280 |
| sctagger | ScTagger-compatible (10x 3' v3) | barcode | 16 | 10 | 10x_3v3 | 6,794,880 |
| splitseq | Parse Biosciences SPLiT-seq / LR-SPLIT-Seq | barcode_1, barcode_2, barcode_3 | 8,<br>8,<br>8 | 10 | splitseq_bc2 (bc1), splitseq_bc1 (bc2, bc3) | 96 per position |
| visium | 10x Genomics Visium Spatial Gene Expression | barcode | 16 | 10 | 10x_Vis_V1 | 4,992 |
| visium_hd | 10x Genomics Visium HD Spatial Gene Expression | barcode_1, barcode_2 | 15–16, 14–15 | 9 | visium_hd_bc1, visium_hd_bc2 | 3,350 each |
| nanopore_rapid_bc | ONT Rapid Barcoding Kit (bulk) | none<br>linker-embedded) | N/A | N/A | N/A | N/A |

Each row corresponds to a named layout key accepted by the -l / --layout flag. Whitelist sizes reflect the default kit alias used in each layout CSV; additional kit aliases for other chemistry versions are supported (e.g., 10x_3v1–10x_3v4, 10x_5v1–10x_5v3, 10x_Vis_V1–10x_Vis_V5) and can be specified by editing the whitelist field in a custom layout CSV. bc: barcode; N/A: not applicable.

### Broad, rapid, and efficient LRS data analysis

To showcase RAD’s performance on a cutting-edge data modality, we first applied RAD to a publicly available 10x Visium HD long-read dataset of murine brain parenchyma^18^. This dataset comprises a single tissue slice sequenced across four Nanopore PromethION flow cells, returning an average of 150M unprocessed reads per replicate and a total of 600M reads in aggregate. We compared per-spot log normalized counts produced by the standard workflow, Epi2ME’s Percula^19^, to those recovered by RAD during whitelist generation (**Figure 1B**) on one technical replicate using SpaceRanger’s *in_tissue* annotation as reference for valid spots. Across the approximately 120K spots in tissue, RAD detected 45.5M reads with no barcode errors compared to Percula’s 36.6M million and averaged 25% more mean reads per spot (369 vs 296). This analysis highlights the potential of LRS to support discovery in new biological applications given accessible computational tools to demultiplex and further analyze the resulting data.

Major goals in the development of RAD were to ensure that it is computationally lightweight and adaptable to cutting edge long-read sequencing datasets. To benchmark computational efficiency, RAD was compared against gold-standard tools Flexiplex and BLAZE on two tasks: whitelist generation and demultiplexing on two individual components of the LongBench dataset^11^ (**Figure 1C, D**). Each tool was run in triplicate with 4, 16 and 64 threads on the single-nuclei Oxford Nanopore Technologies-sequenced (ONT, 119M reads), and single-cell Pacific Biosciences-sequenced (PB, 195M reads) datasets. Using 4 threads, RAD completed processing for single nuclei-ONT on average in under 4.5 hours, and single cell-PB in about 3.5 hours. At the same thread count, BLAZE and Flexiplex ran on average for 33 (10x as long as RAD) and 6-12 hours (1.5-4x as long as RAD). These comparisons excluded the time required to trim residual adapter sequences from BLAZE and Flexiplex outputs with *cutadapt*^20^, a step RAD performs by default.

All three tools utilized under 2.5 GB of total memory in total across all three threading conditions; RAD had the highest memory demands of any tool at 16 and 64 threads due to its use of a custom data structure that contains the entire inclusion list in memory during demultiplexing. Altogether, we found RAD to be the fastest tool, comparable in memory utilization, and uniquely capable of processing cutting-edge long-read spatial transcriptomic sequencing compared to both BLAZE and Flexiplex.

### Performance on synthetic data

RAD’s performance for both whitelist generation and subsequent demultiplexing was benchmarked against BLAZE^17^ and Flexiplex^15^ on the previously published synthetic ScTagger-L dataset^21^ consisting of 5 million reads derived from 5,000 ground-truth barcodes, with errors introduced using the Badread^22^ Nanopore2023 model (**Figure 2A**). For whitelist generation, RAD’s default settings and BLAZE run with the *expected-cells* parameter set to 5,000, both returned all 5,000 true positives while BLAZE returned an additional 15 false positives. Flexiplex, run with the flanking adapter edit distance set to 8, captured only 3,004 true positives but no false positives (**Figure 2B**). Therefore, RAD achieved perfect sensitivity and precision. F1 scores for RAD and BLAZE were nearly identical and exceeded that of Flexiplex. The small difference between RAD and BLAZE could be attributed to the small number of BLAZE false positives, while Flexiplex performance suffered due to its lower sensitivity on these synthetic data (**Figure 2C**).

**Figure 2.**
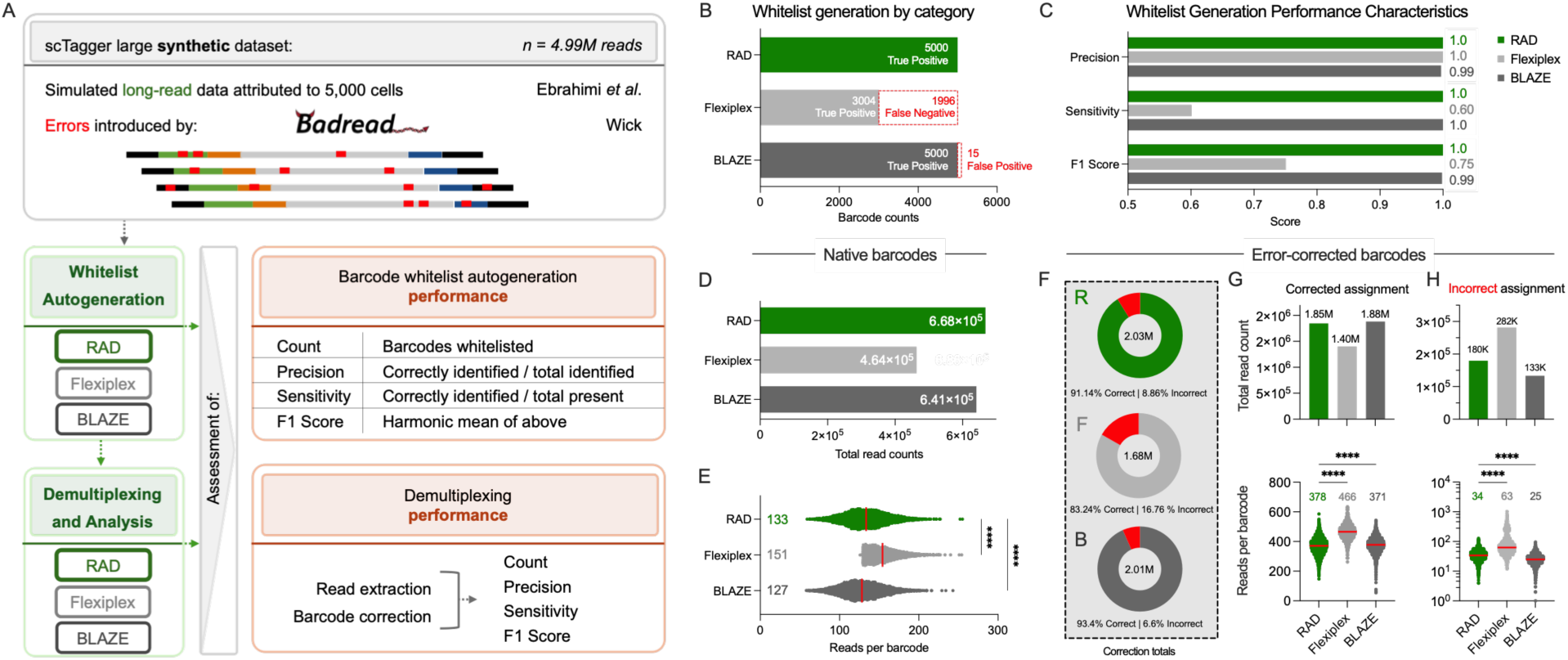
Benchmarking performance on synthetic data. **A.** Schematic of synthetic data used in performance evaluation and comparison of RAD (R, green), Flexiplex (F, light gray), and BLAZE (B, dark gray) in whitelist generation, demultiplexing, and analysis tasks. **B-C**. Whitelist generation performance. Quantitation of the true positive, false positive, and false negative barcodes generated by each method (**B**), and their resulting precision, sensitivity and F1 scores (**C**). **D-E**. Demultiplexing performance on native barcodes. Total read count (**D**) and read depth (**E**) captured for native barcodes by each tool. **F-H**. Demultiplexing performance on error-corrected barcodes. The number of correctly and incorrectly assigned barcodes were quantified (**F**). For both correctly assigned (**G**) and incorrectly assigned (**H**) barcodes, total read counts (top) and read depth (bottom) captured by each tool. Statistical significance of each tool compared to RAD determined by one-way Kruskal-Wallace ANOVA with Dunn’s Multiple Hypothesis Correction (****p < 0.0001). Red bar and numbers in inset indicate median values.

Each tool’s demultiplexing outcomes were examined next, beginning with reads that already exactly matched a true barcode (no correction required) (**Figure 2D**). Here RAD outperformed both tools, recovering about 5% (27K) more reads than BLAZE and 40% (200K) more than Flexiplex, whose performance was significantly compromised based on low sensitivity in its whitelist generation step. Offsetting this limitation, Flexiplex yielded greater mean read depth on true positive exactly matching barcodes, while counts per cell were more similar in magnitude for RAD and BLAZE (**Figure 2E**).

All three tools also error-corrected barcodes to capture more reads from the raw data. Through these processes, RAD and BLAZE rescued more than 2M reads and assigned more than 90% of these reads to the correct barcode, while Flexiplex rescued close to 1.7M reads and assigned about 83% to the correct barcode (**Figure 2F**). RAD attempted correction on more reads (**Figure 2F**) and properly corrected the greatest number, though not proportion, of reads compared to both other tools (**Figure 2G**). All three tools also improperly corrected reads to true barcodes (**Figure 2H**), with Flexiplex incorrectly assigning up to twice as many corrected reads per cell, and in total, than either RAD or BLAZE. In this evaluation, BLAZE achieved the fewest overcorrections, both in terms of number of reads and reads per barcode, closely followed by RAD.

Of the total corrected reads uniquely output by BLAZE and Flexiplex, a proportion (∼60-70K reads, roughly 3% of the corrected total) contained ground-truth barcodes for which Badread introduced errors resulting in exact matches to barcodes from the complete chemistry whitelist that were then corrected to a barcode within the tool’s shortlist. These errors in demultiplexing did not extend to RAD, which was designed with explicit guards against these occurrences. Corrections that generate these errors are disallowed in standard short-read sequence demultiplexing^23^, as they can lead to significant downstream computational artifacts^24–26^. This observation prompted the comparison of RAD, BLAZE, and Flexiplex’s abilities to protect against various false-positive barcode correction errors across multiple real data types.

### Avoidance of common demultiplexing errors

To more directly compare rates of overcorrection by these tools, the propensity of each to assign reads to false positive barcodes was quantified by supplying each tool with a fixed whitelist, overriding the automated whitelist generation step, comprising a size-matched set of barcodes not among the ground truth or matched short-read data from synthetic data, gold-standard real-world data, and specialized targeted long-read single-cell (TLR-SC) RAGE-seq^27^ data that used hybridized sequence probes to pull down immune-receptor sequence isoforms (**Figure 3A**). In this context, the majority of reads were expected to be assigned through errant correction rather than by direct matching, and indeed, the vast majority (>98%) of total reads returned for all tools could be attributed to overcorrection. Across analytical replicates for all three datasets tested, Flexiplex and BLAZE compiled more false positive reads and at greater read depth per barcode than RAD (**Figure 3B,C**). Consistent with its requirement of sufficient exactly matching barcode support before assigned corrections are accrued that no more than 10 reads be error-corrected to a given barcode without observation of at least one exact (uncorrected) match, RAD generated fewer than 50K false-positive read assignments; BLAZE and Flexiplex assigned up to 30-fold as many total false positives (**Figure 3B**). Per barcode, RAD assigned significantly fewer false positive reads than either BLAZE or Flexiplex (**Figure 3C**). Taken altogether, RAD exhibited substantially fewer false-positives in these overcorrection tests than comparator tools.

**Figure 3.**
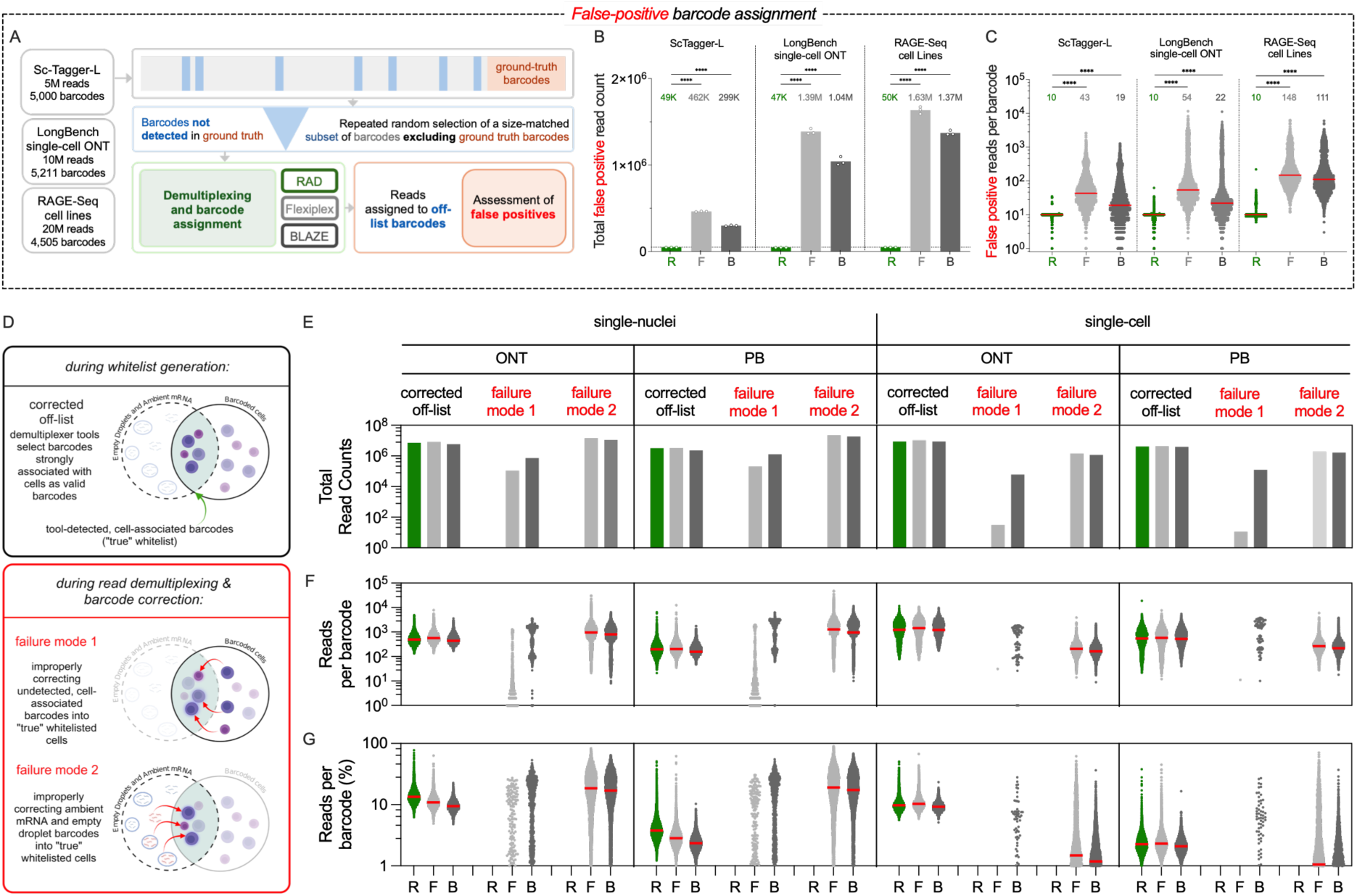
Adaptive whitelisting provides high fidelity barcode identification and correction. **A-C**. Characterization of false positive barcode assignment rate. Using three datasets, random, size matched sets of “off-list” barcodes were provided as the whitelist to each tool and the rate at which reads were assigned to this list was assessed (**A**). Quantification of the of total read count (**B**) and read depth (**C**) assigned to false positive barcodes by each tool. Statistical significance determined by one-way ANOVA (****p < 0.0001). **D-G**. Barcode correction outcomes (**D**). Using the LongBench dataset, the total read counts (**E**), the read depth (**F**), and the percentage of reads per barcode (**G**) derived from corrected off-list or the two failure modes were quantified for each tool. Red bar and, where present, numbers in inset indicate median values.

Recognizing that this test represents a worst-case scenario, we next sought to explore the occurrence of improper barcode correction in real datasets. To conduct this analysis, the LongBench datasets purpose-built for benchmarking the sensitivity and precision of LRS tools, were utilized. We processed each of the four LRS LongBench datasets with RAD, BLAZE and Flexiplex and characterized barcode corrections, including errors classified into two modes (**Figure 3D**). While extracted barcodes not originating from the CellRanger inclusion list and corrected to an entry in the generated whitelist were considered properly assigned and enumerated as “corrected off-list”, failure modes comprised improperly corrected barcodes from the inclusion list. Failure mode 1 was defined by reads from barcoded cells missed during whitelist generation that were corrected into another barcode, and failure mode 2 was defined as reads unaffiliated with either the long-or-short-read selected whitelists and thus considered to represent ambient background barcodes that were corrected into a tool-whitelisted barcode.

Whereas all three tools assigned reads from “corrected off-list” barcodes with similarly high read counts and read depth per barcode, their correction failure mode profiles differed considerably (**Figure 3E-G**). BLAZE and Flexiplex assigned reads via the two failure modes, contributing a substantial percentage of the reads per barcode (**Figure 3G**) and could be detected in samples generated from both single nuclei and cells, as well as in data generated using PacBio (PB) and ONT technologies. In most samples tested, Flexiplex was less prone to mode 1 than BLAZE (**Figure 3E, F**). For both BLAZE and Flexiplex, the reads generated by mode 2 occurred at a read depth comparable to that of the corrected off-list read depth for all three tools (**Figure 3F**). These errors were particularly abundant in the single-nuclei processing, outpacing properly corrected reads per cell by nearly two-fold. Overall, these overcorrection experiments highlight the value of RAD’s adaptive whitelist generation, which provides high fidelity demultiplexing of reads while also rescuing error-affected reads with comparable performance to existing tools.

### Comparative benchmarking on 37 long-read datasets on whitelist generation

All three tools were benchmarked across a breadth of long- and short-read paired sequencing data including 12 single-nucleus^11,28^, 11 single-cell ^4,11,17,29–31^ 9 Visium^32,33^, 3 targeted long-read^27,29^ and 2 synthetic^21^ samples, for 37 long-read samples in total (**Figure 4A**). These datasets ranged across short-read whitelist sizes, total read counts, GC content, and read quality (average Q-score) (**Figure 4B-E**). Each tool was used to generate whitelists for each long-read sample, and F1 scores, precision, and sensitivity were quantified using paired short-read sequencing whitelists as ground truth (**Figure 4F-H**). Overall, RAD achieved a significantly higher F1 across the 37 samples compared to Flexiplex and BLAZE, yielding the highest F1 scores on 26 of the 37 datasets (70%) (**Figure 4I**). Differential scores were driven primarily by a trade-off between precision and sensitivity; while BLAZE and Flexiplex were more precise (**Figure 4J**), RAD recalled more valid barcodes (**Figure 4K**).

**Figure 4.**
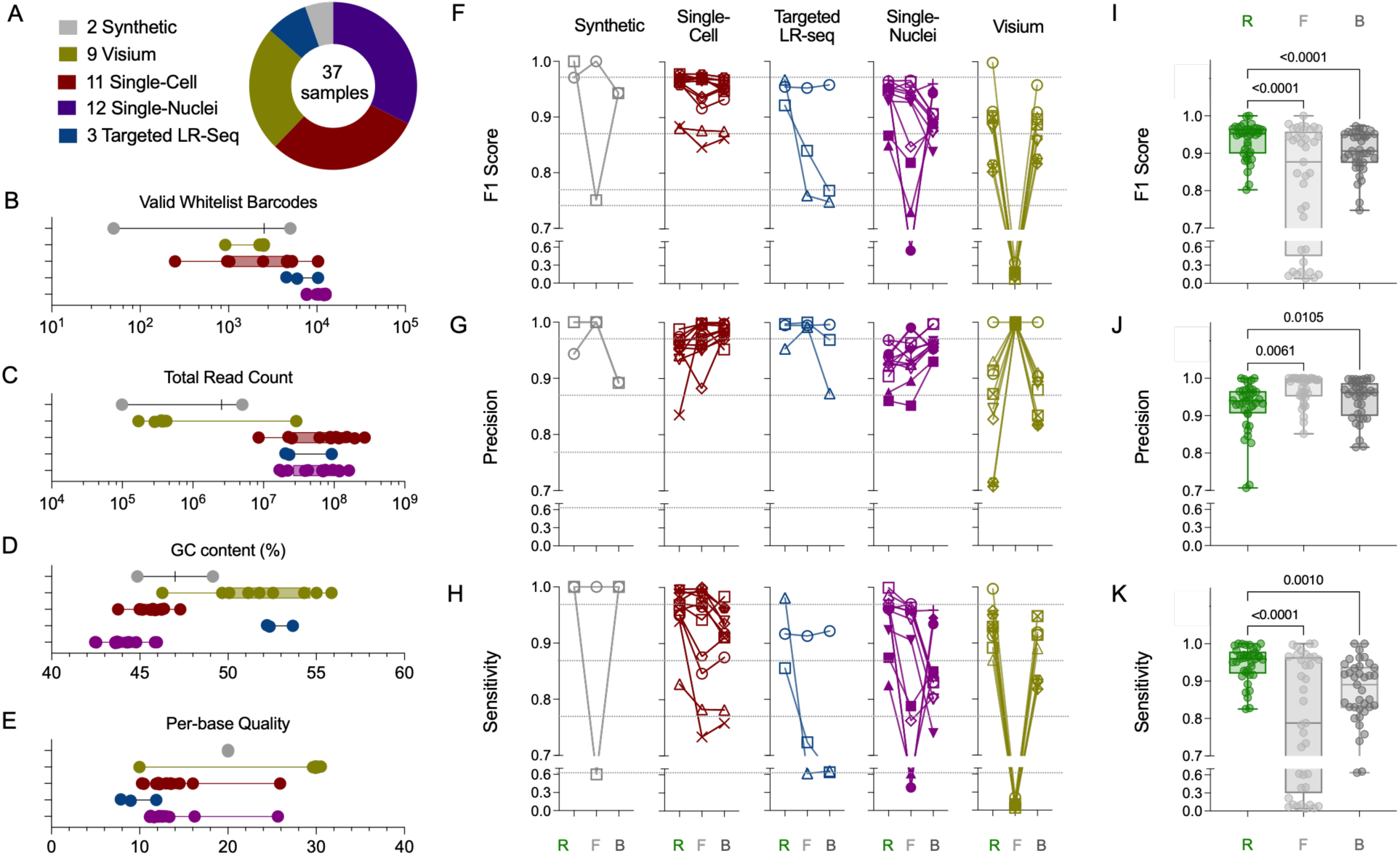
High performance across diverse datasets and read architectures. Long-read data generated from 37 samples were chosen from a variety of experimental protocols, sample types, and read architectures (**A**). The short-read data characteristics have been plotted to illustrate the diversity of the underlying data **B-E.** This characterization included valid whitelist barcodes (**B**), total read count (**C**), GC content (**D**), and average Q (**E**). The long-read demultiplexing performance of each tool using RAD (R, green), Flexiplex (F, light grey), and BLAZE (B, dark grey), was characterized using F1 scores (**F**), precision (**G**), and recall (**H**) for all 37 samples. The distributions and medians of F1 scores (**I**), precision (**J**), and recall (**K**) across all 37 samples are shown as box-and-whisker plots, with whiskers indicating the minimum and maximum values. Statistical significance determined by Friedman Test.

### Robust handling of targeted and complex LRS datasets

Beyond the broad applicability of RAD, a central consideration of its development was to make demultiplexing of targeted sequencing and complex read structures straightforward and robust to error at both the whitelist generation and demultiplexing stages. The RAGE-Seq long-read dataset^27^ comprises roughly 20 million reads captured by targeted exon-probe hybridization and amplification, with paired short-read data from a mixture of Ramos (B-cell) and Jurkat (T-cell) lines and PBMC-derived monocytes (**Figure 5A**). Demultiplexing of these data allows for assessment of both barcode assignment and accurate mapping of immune receptors by comparison to the short-read defined barcode and receptor identification. Compared to Flexiplex and BLAZE, RAD detected more cells in total (**Figure 5B**) and of each major cell line type (**Figure 5C**). RAD also captured more receptor reads than the other two tools for both Ramos and Jurkat cells (**Figure 5D**). Much of this increased read quantity came from robust and faithful correction of receptor reads (**Figure 5E**). RAD’s detection of reads and cells translated into slightly higher read depth for the Ramos cells compared to the other tools (**Figure 5F**). While Flexiplex had higher read depth for the Jurkat cells, Flexiplex missed 22% of the Jurkat cells that RAD detected (**Figure 5F**).

**Figure 5.**
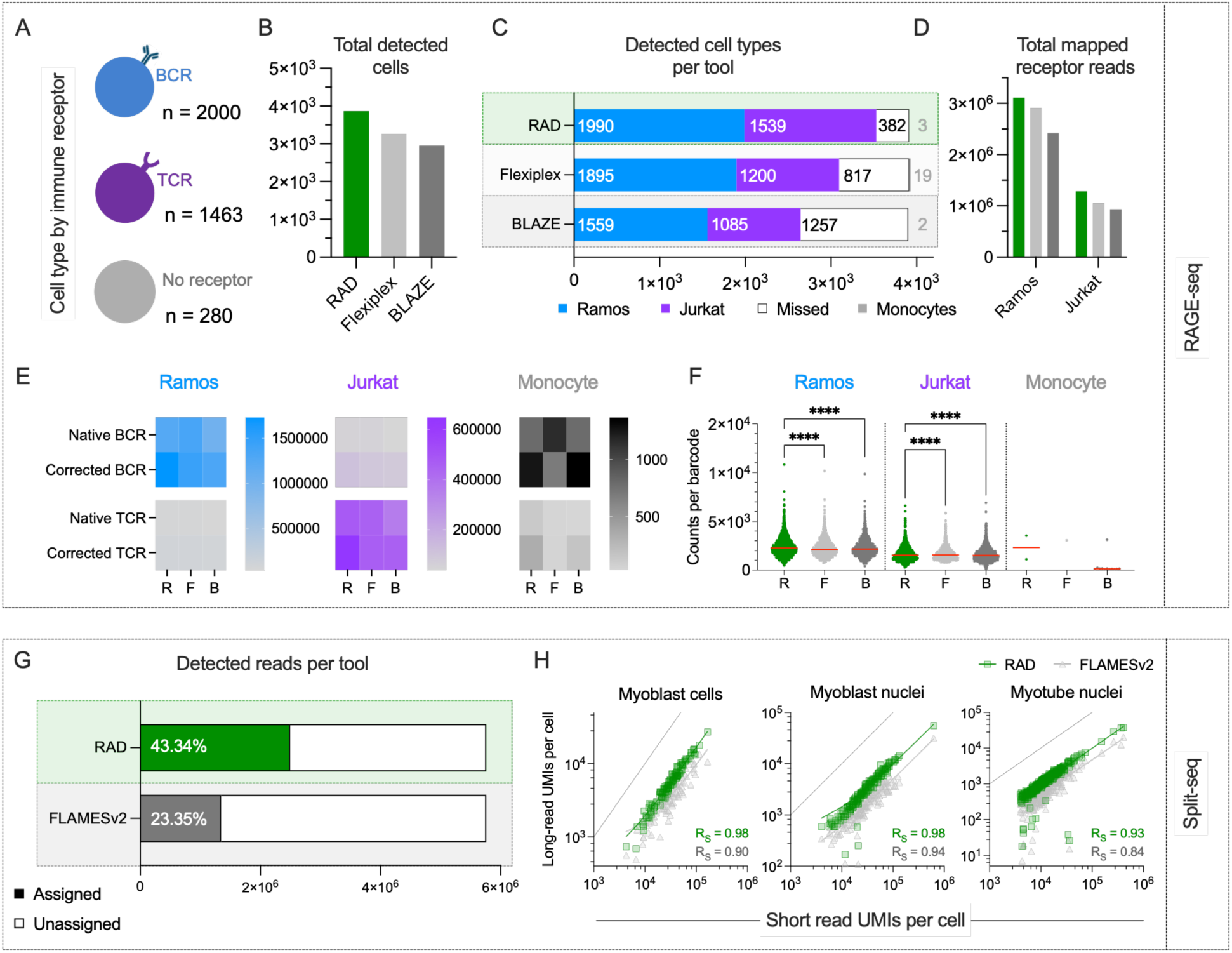
Benchmarking RAD performance on data with complex read structures. **A-F**. Performance of each tool in demultiplexing and mapping RAGE-seq-derived immune cell receptor reads when applied to a mixed pool of immune cells with distinct receptors. Composition of input cells (**A**). Comparison of total cells detected by RAD (R, green), Flexiplex (F, light gray), and BLAZE (B, dark gray) (**B**), number and identity of cell of each type (**C**), and the number of BCR and TCR reads mapped (**D**) by each tool. Number of reads with native or corrected barcodes (**E**) and read counts per barcode (**F**) mapped to each cell population by each tool. Statistical significance by paired two-tailed t-test (****p<0.0001). **G-H**. Performance of RAD (green) and FLAMESv2 (gray) in demultiplexing LR-Split-seq data. Detection of barcodes in LR-Split-seq dataset in which assignment requires successful identical of three barcodes per read (**G**). Long read and short-read UMI counts per cell were compared across the three cell types annotated in the original paper per tool (**H**). Dotted lines represent identity (x = y); Spearman correlation coefficients (R_S_) are presented in inset along with best fit lines for each tool.

A critical shortcoming of LRS is that sequencing errors make UMI-based quantification of molecules difficult^34^. Further, the complex triple barcoding system of Split-seq read architecture means that many long-read demultiplexers are unable to capture all three barcodes as well as UMI sequence. FLAMESv2^35^ was developed to be highly configurable to these complex applications of LRS data and to quantify UMI. To test RAD’s performance on a data type with complex barcoded structures, both tools were compared on a LR-Split-seq dataset^8^ that employed split-pool combinatorial barcoding. RAD detected nearly twice as many reads as FLAMESv2 from this dataset (**Figure 5G**). Reads were recovered for all 568 barcodes across the two myoblast (cells and nuclei) and single myotube nuclei samples detected in the dataset’s paired short-read sequencing. Per-cell UMI abundances correlated strongly between long- and short-read sequencing, though RAD (mean R_S_ = 0.98) captured more UMIs and better preserved relative per-cell UMI abundance as compared to FLAMESv2 (mean R_S_ = 0.89) across all three samples (**Figure 5H**). Taken together these results suggest that RAD may be an enabling tool for demultiplexing LRS data, particularly in cases of complex read architecture or when UMI quantification would be useful for exploration of expression data.

### Improved annotation and data depth

To test how demultiplexer tool choice can affect usable cell identities and downstream differential gene (DEG) analysis, RAD, Flexiplex, and BLAZE were applied to the single-nucleus ONT LongBench data^11^ comprising a mixture of eight cell lines derived from three different lung cancer types: adenocarcinoma (LUAD), and small-cell lung cancer types -A and -P (SCLC-A and SCLC-P, respectively). Here, genomic variation between cell lines detected in the bulk short-read data was used to annotate the single-nuclei data as previously reported^11^, and graphically presented in transcriptomic space as defined by Uniform Manifold Approximation Projection (UMAP), providing a ground truth comparator to the demultiplexing and cell line annotation of nuclei using the long-read data (**Figure 6A**). Nuclei without adequate short-read data coverage of the genomic variants used as markers of cell line identity were denoted as short-read un-annotated and served as a basis for additional biological information and recovery.

**Figure 6.**
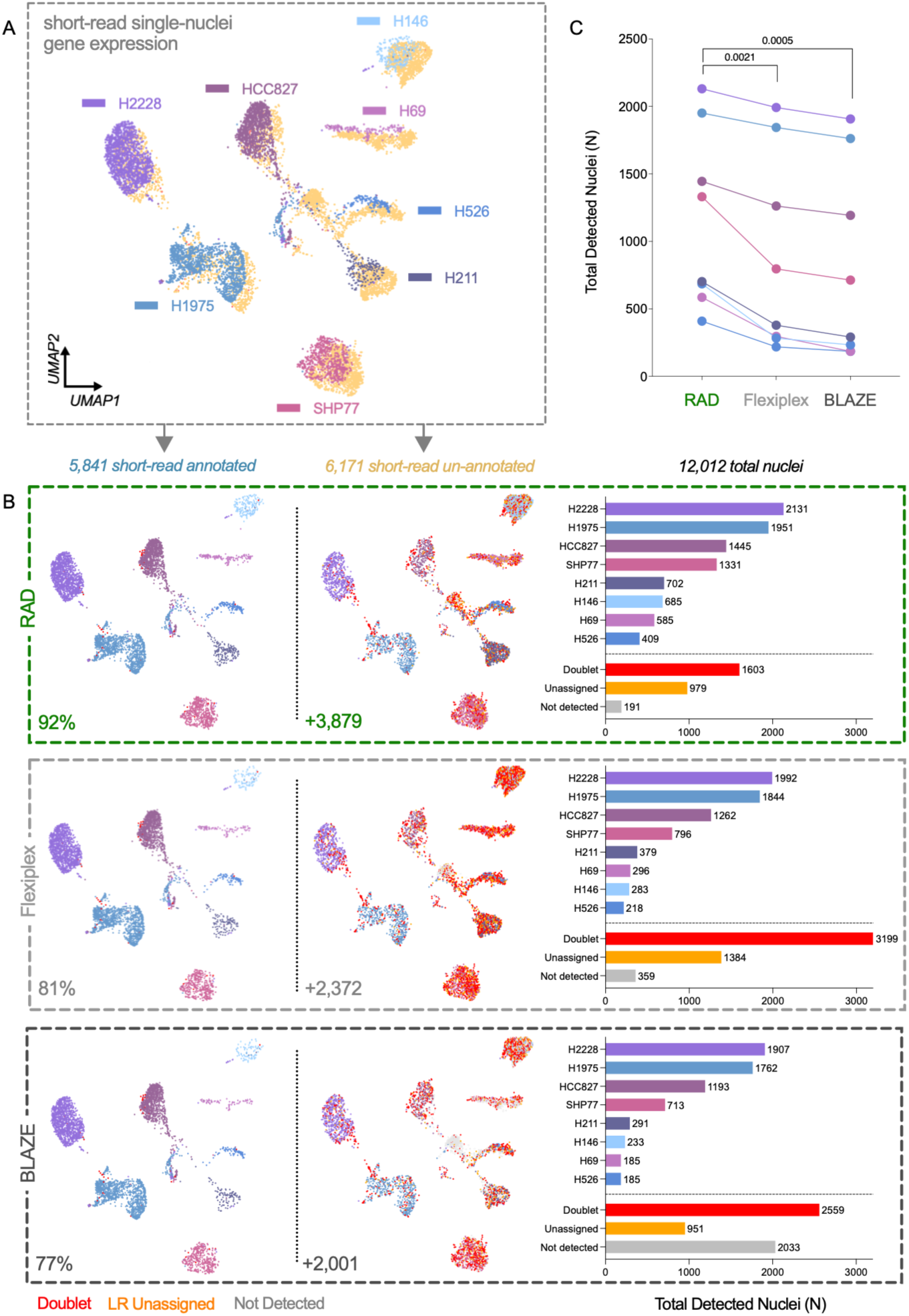
Improved single-nuclei data depth and cell line annotation. **A**. Uniform Manifold Approximation Projection (UMAP) plot of LongBench single-nuclei dataset depicting short-read un-annotated (peach, offset for visualization), and short-read annotated (other colors) nuclei per cell line. **B**. Performance of each tool in identifying short read-annotated (left) and short read-un-annotated (center) nuclei on UMAP axes defined in A, with quantification of each cell line or annotation status (right). Values in inset represent the percent (left) and number (center) of tool-annotated nuclei. **C**. Total number of nuclei identified by each tool per cell line. Statistical significance defined by one-way ANOVA with Dunnett’s multiple hypothesis correction.

The concordance between short- and long-read tool annotations and transcriptomic profiles across all three tools was measured (**Figure 6B**). After demultiplexing, the number of nuclei assigned to each cell line was quantified, along with those containing variants from more than one cell line that were classified as doublets, and those that were unassigned to a cell line type or that were not detected in the long-read data. More of the roughly 6K short read-assigned nuclei were correctly detected and assigned by RAD than by either Flexiplex or BLAZE. Additionally, among the similar number of short read un-annotated nuclei, RAD also annotated a greater number than comparator tools, a feature most apparent in nuclei with high UMAP dimension 1 values. Most of the nuclei that could not be assigned to a cell type based on long-read demultiplexing results were doublets, which presented a greater proportion (nearly 30%) of reads with errant correction (failure mode 2) as compared to singlet-assigned nuclei (13%). Overall, RAD detected more nuclei than did Flexiplex or BLAZE (**Figure 6C**), returning nearly twice as many nuclei of the sparsest cell lines compared to either BLAZE or Flexiplex (**Figure 6C**).

### Enhanced biological discovery

Finally, to determine RAD’s added biological value, two sparsely annotated cell lines or different cancer subtypes (SCLC-A, H146, and SCLC-P, H526) were chosen to test the effect of demultiplexing tool performance on DEG analysis. Briefly, the paired short-read gene expression data was utilized as a ground truth reference in analysis restricted to the genes detected across all three long-read demultiplexers and associated short-read data. By demultiplexing the data using RAD, more genes were identified as differentially expressed between the two cell lines (**Figure 7A**). Of the 1,181 DEG (adjusted p < 0.05 and two-fold change) that were defined by short-read data, RAD recovered 654, compared to 556 for Flexiplex and 522 for BLAZE. Of these, five times more reference-supported DEGs were uniquely detected by RAD. As a result, RAD was most sensitive and comparably precise to the other tools, yielding the highest F1 score **(Figure 7A**).

**Figure 7.**
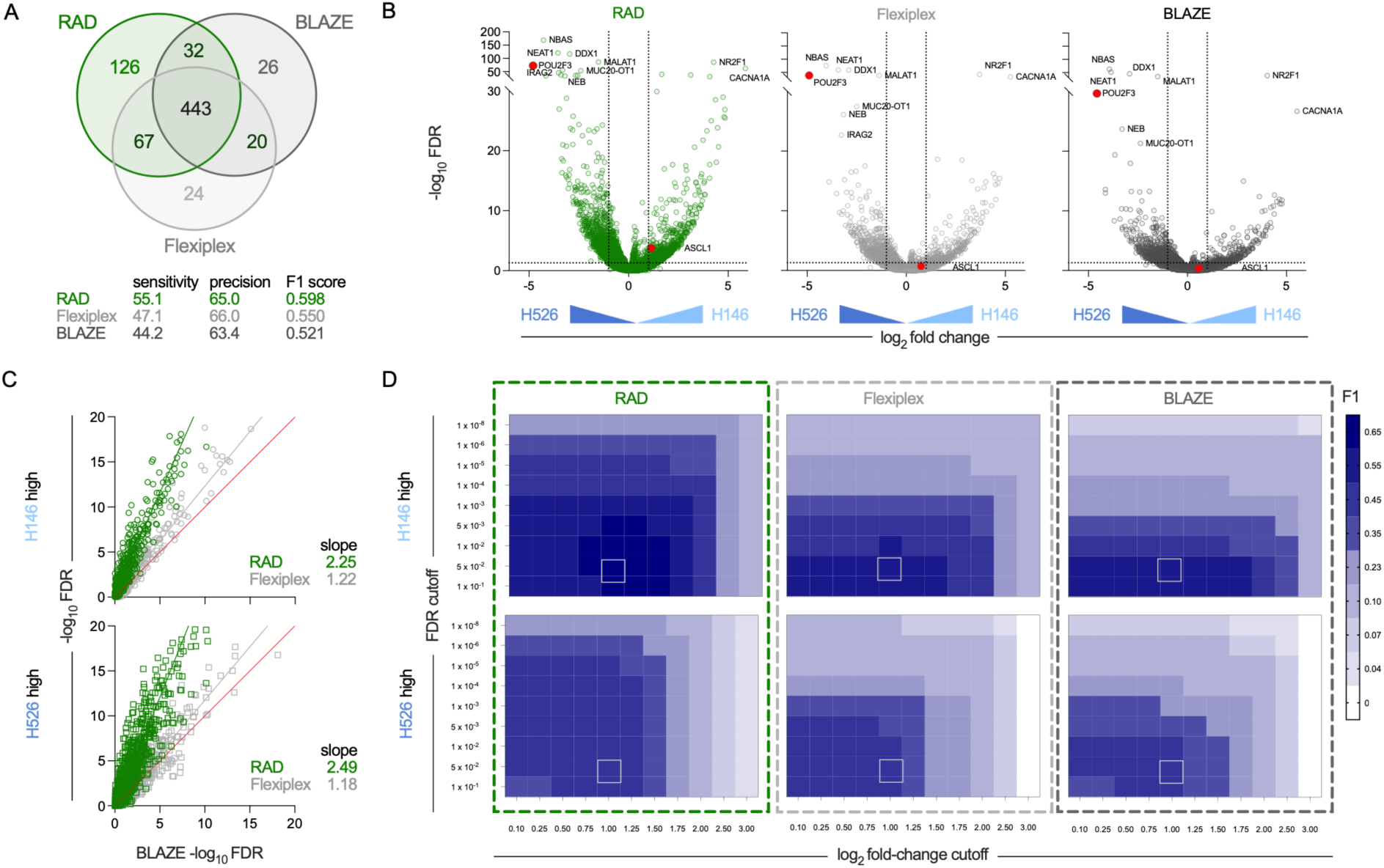
Faithful and confident identification of differentially-expressed genes. **A**. Venn diagram depicting the overlap between genes identified as differentially expressed (FDR > 0.05, fold-change >2) by each tool. Tool sensitivity, precision, and F1 scores for identification of genes defined as differentially expressed in short-read data. **B**. Volcano plot depicting the fold change and confidence (FDR, False Discovery Rate) of differences in gene expression between H526 and H146 cell lines. Genes identified with greatest confidence (FDR > 10^−20^) in Flexiplex or BLAZE are identified by name. Genes previously identified as key transcriptional regulators and markers of small cell lung cancer (SCLC) types -A and -P, ASCL1 and POU2F3, are indicated in red. Dotted vertical and horizontal lines indicate two-fold expression changes and FDR of 0.05, respectively. **C**. Scatterplot of confidence in expression differences between BLAZE (x-axis) other tools (y-axis) for genes that are overexpressed in H146 (top) and H526 (bottom) nuclei, respectively. Red line indicates unity and slopes of best fit lines for RAD and Flexiplex are listed in inset. **D**. Accuracy of differentially expressed gene identification for each tool (F1 scores) as compared to ground truth, as defined by short read data across a sweep of fold-change (x-axis) and FDR (y-axis) cutoff thresholds. Highlighted box indicates the condition presented in panels A-C.

Importantly, RAD was the only tool to identify both key transcriptional regulators used to categorize SCLC cell types, POUF2F3 (SLCL-P)^36,37^ and ASCL1 (SCLC-A)^37,38^ as DEG in these data (**Figure 7B**). One clear benefit of demultiplexing using RAD was the increased confidence in expression differences. By comparing false discovery rates on a gene-by-gene basis, data processed using RAD identified differential expression with about twice the confidence of the other tools (**Figure 7C**). Using the short-read expression as ground truth, the accuracy of the differentially expressed gene analysis could be assessed for each tool by F1 score. Across both false discovery rate (FDR) regimes and degrees of fold change, RAD consistently classified genes as differentially expressed with better precision and sensitivity to the short-read data than the other tools (**Figure 7D**). Of the 99 FDR and fold change threshold combinations evaluated, RAD achieved the highest F1 score in 90% of test cases (**Figure 7D**). Altogether, these results demonstrate RAD’s significantly better recovery of reference-supported DEG signal, and better sensitivity-precision balance compared to competitor tools across the majority of evaluated thresholds. This analysis demonstrates how single cell long read sequencing data can be useful not just for better understanding of isoforms and gene architecture, but also for sensitive detection of differential gene expression to support biological discovery.

## Discussion

Existing tools, including BLAZE^17^ and Flexiplex^15^, have expanded long-read demultiplexing capabilities, but remain limited by one or more meaningful constraints: rigid read-structure assumptions, benefitting from short-read sequencing for optimal results, and challenges in addressing complex long-read-specific error modes. To address these constraints, we developed RAD, a read-structure agnostic demultiplexer for LRS. RAD includes pre-configured read layouts and reference whitelists covering 10x 3’/5’, Visium, Visium HD, LR-Split-seq, and Nanopore bulk chemistries.

Across 37 datasets, RAD discovered barcode whitelists with higher precision and sensitivity than existing tools, generated substantially fewer false-positive reads than the next-best tool in an orthogonal overcorrection test, and completed whitelist generation and demultiplexing on the LongBench datasets in a fraction of the time required by Flexiplex or BLAZE while maintaining comparable memory utilization. These advantages underscore RAD’s capacity to be first-in-class at detecting barcodes, demultiplexing LRS, and returning high-fidelity results driving biological discovery.

Another compelling application of RAD is to TLR-SC of the highly variable B- and T-cell immune receptors (BCRs and TCRs) that recognize specific antigens to trigger adaptive immunity^39,40^. As many single-cell methods are only capable of sequencing the 3’ or 5’ ends of the transcript, they either do not capture the sequences that define receptor identity (3’ limitation), or miss class switching information important for B cell biology^41^ (5’ limitation). RAD is first-in-class at demultiplexing complex experimental formats such as RAGE-seq, which can be applied to overcome these short-read limitations and return valuable information on receptor state and identity throughout immune development.

RAD includes algorithmic approaches to address current limitations. To accept arbitrary read structures, RAD parses a user-defined comma-separated value (.csv) file to lay out static (adapters, linkers, primers, poly-T tails) and variable (barcodes, UMIs, reads) elements from 5’ to 3’. To reduce, or arguably eliminate, short-read dependence, barcode whitelists are derived *de novo* based on deriving a data-driven cell-background boundary rather than from an external reference. Together, read-structure agnosticism, unpaired operation, and robust error handling position RAD for current and future sequencing modalities for which standard demultiplexing pipelines do not yet exist.

RAD is agnostic to sequencing layouts, but not to prior knowledge of their content: users must still supply the sequences of static elements (adapters, linkers, primers) in the layout CSV. RAD’s whitelist generation has not completely matched that unification; while a single layout engine can accept arbitrary single-cell and spatial read structures, RAD’s algorithm demarcating cell from background in 10x-style single cell libraries does not currently generalize to Visium HD. This limitation requires a distinct whitelist generation solution rather than an adjustment of the pre-existing algorithm. This exception notwithstanding, we expect that as long-read single-cell and spatial protocols continue to diverge from 10x-anchored chemistries, RAD’s layout abstraction can concurrently extend without tool re-engineering. A subsequent reformat step writes spatial coordinates into read headers, collapsing a typical multi-tool coordinate-mapping pipeline into a single unified step.

We anticipate near-term applications in MAS-Iso-Seq and single-cell Iso-Seq, which concatenate transcripts for deeper isoform coverage^9^; we hope to expand this deconcatenation approach to R2C2-style reads. Another priority is UMI correction: RAD currently validates UMI hits through positional extraction but principled UMI correction under long-read error profiles remains an open problem across current tools. Extending RAD’s dual-bookkeeping approach from the barcode layer to UMI correction is a natural next step toward clean long-read count matrices.

Beyond core demultiplexing, RAD’s flexible output can be easily processed by a variety of custom downstream analysis pipelines. RAD will be particularly valuable for groups without dedicated computational infrastructure, for whom a single installable pipeline, rather than a chain of coordinated dependencies, lowers the barrier to long-read single-cell data analysis.

## Methods

### Algorithm Overview

#### Read Structure Annotation

RAD takes as input a read layout .csv file that defines the structure of a raw read from 5’ to 3’ in terms of static and variable elements. Static elements have known sequences: for example, primers, adapters, and linkers. RAD also handles special static elements, such as poly-T tails, for which the user knows the sequence, but the length is undefined. Variable elements are those with unknown sequence within the read; this group is constituted of captured reads, UMIs, and barcodes. For user convenience, RAD ships read layout template files for several common sequencing modalities: 10x 3’ Chromium, 10x 5’ Chromium, 10x Visium, 10x Visium HD, LR-Split-seq, ScTagger-style synthetic, and Nanopore rapid barcode (**Table 1**). Custom read layouts can also be passed as a file path. Because RAD will parse a read in both forward and reverse-complement directions, RAD will automatically generate a reverse-complement read layout annotation internally based on the user-specified read layout annotation. This feature lets RAD determine the optimal read direction along with sequence ranges for each element.

#### Identifying static elements and inferring mapping parameters for variable elements

RAD first determines the position range for static elements with unknown exact positions, such as adapters, which can serve as anchors for finding variable elements downstream. For many reads, static elements will be different from their pre-defined sequences due to sequencing errors. To robustly identify the static elements for such reads, RAD heuristically determines static element position ranges and allowable sequencing errors through reads in which all static elements in one direction align with no errors (all edit distances of 0). The position range for a static element can be directly derived from these perfect reads. To determine the allowable sequencing errors for a static element, RAD aligns each static element against the perfect reads in the opposite direction. This opposing alignment provides the edit distance thresholds to avoid inferring the wrong strand reads with imperfect static elements. RAD then tabulates the summary statistics of the start and stop positions as well as the edit distances of these misalignments. Per-adapter edit-distance thresholds are derived from the observed mean and standard deviation of this distribution, allowing RAD to adapt stringency to each dataset’s error profile rather than relying on a single global tolerance. These summary statistics can be collectively termed as the misalignment thresholds.

RAD utilizes the arrangement of static elements to derive relative start and stop positions for variable elements. This process can be roughly described with the read layout structured as a series of horizontal elements where “left/upstream” and “right/downstream” describe the 5’ and 3’ direction, respectively. First, RAD uses the generated layout to find the element with a user-predefined read class annotation. It then splits all elements as left or right of the read. RAD then maps the nearest adjacent static sequence for variable elements to the left or right. For example, the static element immediately to the left of a barcode in a standard 10x 3’ sequencing experiment is a TruSeq Read 1 adapter (“forward primer”) with a known sequence and length. As a result, the barcode start position can then be indexed as the stop position of the forward primer, with a single base offset (+1). The barcode stop position is indexed as the stop position of the forward primer with a single base offset (+1), plus the expected length of the variable element (+16, in the case of a standard 10x barcode).

This process is repeated for all variable elements to the left of the read, iterating from left to right. The process is then repeated for all variable elements to the right of the read, iterating from right to left. Once all other variable reads are annotated, the element corresponding to the read is annotated. The read is annotated by finding its immediate upstream and downstream elements and labeling the start and stop positions accordingly. For example, in the same 10x 3’ sequencing layout, the read start would be defined as the stop position of the poly-tail with a single base offset (+1). The read stop would be defined as the start position of the template-switch oligo (TSO) sequence, with a single base offset (−1).

RAD generates primary position maps, as described above, as well as secondary position maps, for all variable elements, so as to account for errors that occur in typical long-read sequencing experiments. For example, some reads may not contain a poly-tail sequence or may be prematurely truncated and lack the reverse adapter. In these cases, RAD utilizes the secondary position mapping of the read element. Here, the secondary read element start is mapped to the stop position of the UMI with a base offset, and secondary read stop is simply mapped to the end of the sequence. This solution allows RAD to gracefully handle prematurely truncated sequences and unreliable static features, such as poly-tails and small error-prone TSOs. The misalignment thresholds and position map are also automatically saved in RAD’s output directory as a single .csv. These files, in conjunction with the read layout file from above, can be modified and reused for subsequent experiments with the same read structure.

### Read Processing & Concatenate Resolution

RAD first generates alignments for all possible static elements in a single sequence in both the forward and reverse directions. Static element positions are found in two passes. RAD utilizes Edlib^42^, configured in local mode with wildcard equalities so that N bases in the static element match any nucleotide to rapidly detect matches in a first pass. Edlib returns one or more candidate intervals with low edit distances for each static element. If the Edlib alignment fails, falling outside the misalignment thresholds or returning multiple ambiguous matches, RAD utilizes the fast striped Smith-Waterman algorithm^43^ to return the location and Compact Idiosyncratic Gapped Alignment Report (CIGAR) string of the optimal alignment match. This two-step process allows RAD to recover truncated static elements, such as terminal adapter sequences, and supports identification of chimeric joins. Detected static elements are masked in place to prevent re-alignment or spurious extraction in downstream variable element processing.

RAD next detects variable elements’ positions using the static element coordinate references generated during position mapping. Like in position mapping, RAD first maps all left-hand (upstream) non-read variable elements, such as barcodes and UMIs, relative to their nearest static anchor, propagating expected start and stop coordinates downstream. It then repeats this process for non-read variable elements downstream in the reverse order. Once all other variable elements have been annotated, RAD computes the boundaries of the read element itself. This procedure ensures that all variable elements have a possible annotation, agnostic to the presence or absence of different static elements. RAD finally applies read-level filtering and concatenate resolution after all possible element positions are annotated. RAD broadly filters reads with structurally inconsistent layouts, missing mandatory variable elements, anomalously short read element lengths, and overlapping elements. RAD also detects and annotates reads as concatenates if both forward and reverse orientation static elements are independently valid in a single read (i.e., each orientation contains correctly ordered, length-consistent static and variable elements that satisfy misalignment thresholds), indicating that two or more complete molecules have been captured in the same sequencing read.

### Barcode Storage Structures

There are variable elements for which the user knows sequences come from a large, predefined set, such as cell barcodes and barcode whitelists. Each barcode is encoded as a custom class *int64_seq*, where each base is encoded as two bits (A = 00, C = 01, T = 10, G = 11). This encoding format allows for maximum barcode lengths of up to 32 bases. For unique entries in any whitelist, *int64_seqs* are stored in a custom class that additionally contains several thread-safe counters for updating barcode counts throughout runtime. Prior to demultiplexing, RAD first loads all sequences in the broader barcode whitelist per unique user-annotated barcode element in the read layout. This architecture supports high-throughput sequencing whitelists and is currently validated on whitelists ranging from roughly one to eleven million barcodes. Larger whitelists are supported in principle. Paired short read data, which usually presents lower sequencing error rates, can serve as a more specific, predefined barcode set for the corresponding long-read data. RAD can define such barcodes as the high-confidence whitelist (typically the CellRanger-filtered whitelist, or a specific subset of cells). If the high-confidence whitelist is between 10-30K entries, RAD can generate and store all barcode variants within a default Levenshtein distance of 2 from each original barcode. Each of these mutated variants is stored linking back to their original barcode(s). Larger “true” whitelists skip this pre-enumeration step and resolve queries through the seed-pigeonhole shortlist described below. RAD also generates a blacklist containing barcode length-sized k-mers of adapter sequences with up to a Levenshtein edit distance of 2.

Computationally, the barcode mapping structure employs a two-level storage structure optimized for both concurrent access and memory efficiency. Unique barcode sequences are stored as an unordered set. The relationship between mutated variants and their corresponding “correct” barcode is stored in a parallel hashmap, in which the key is the mutated variant and the value is a stable pointer to the unique correct barcode sequence. This hashmap is pre-populated at load time using the variants generated above; subsequent demultiplexing hits the cache in constant time. The hashmap structure allows for multiple counters to be concurrently accessed and modified over RAD’s runtime in a thread-safe manner. For example, during the barcode correction algorithm, RAD can store the incorrect variant mapping back to a single correct barcode. Subsequent recurrences of the incorrect variant return the cached mapping in constant time, avoiding repeated full-distance correction. At the top level, each whitelist instance holds two independent tables: *true_bcs* (the high-confidence whitelist, generally the CellRanger-filtered whitelist or a specific subset of cells; optionally expanded into a *filtered_bcs* substructure of mutation-generated variants) and *global_bcs* (a broader set, e.g., the CellRanger barcode inclusion list). Both are indexed independently, and queries fall back from *true_bcs* to *global_bcs* to maximize sensitivity without relaxing correction thresholds.

### Barcode Filtering & Correction

After extraction, barcodes and their reverse-complement sequences are filtered for extraction artifacts: homopolymer runs equal to or greater than 40% of the expected barcode length (minimum four bases), and low-complexity sequences defined as having fewer than four unique 2-mers. Adapter-derived k-mer blacklists (generated up to Levenshtein distance 2 from the flanking static elements in the layout) are also rejected. Barcodes not matching either the broader or high-confidence whitelists enter the correction algorithm. Candidate edit distances are computed with Myers’s bit-parallel algorithm^44^ on the 2-bit-encoded *int64_seq*. To shortlist candidates before full distance computation, RAD builds a seed-pigeonhole index over each whitelist: every barcode is partitioned into three contiguous segments, and each segment is packed into a 32-bit key encoding both the partition identifier and the 2-bit-encoded segment. Keys are stored in an unordered hash map whose values are pointers directly into the whitelist entries.

At query time, the three partitions of the candidate barcode are looked up independently; any whitelist entry within Levenshtein distance 1 of the query must share at least two of the three partitions, so the union of hits across partitions is a superset of all matches within an edit distance of 1 and is typically several-fold smaller than the full whitelist. If a single candidate in the shortlist is within the edit-distance threshold (default: 4), it is accepted only after passing an internal quality gate that compares raw, corrected, and total read counts maintained on each whitelist entry to ensure the candidate has been independently observed in the data. When multiple candidates satisfy the distance constraint, RAD applies a tiebreaker that resolves to the barcode with the lowest edit distance. Ties are broken by higher empirical read support. RAD exposes two correction strategies: *offensive* applies corrections wherever a qualifying candidate exists, maximizing assignment rate, while *defensive* queries the broader whitelist before the high-confidence whitelist and tightens the per-barcode correction budget, reducing false corrections at some cost to recall. RAD also supports both a permissive mode that accepts any passing component barcode, and a strict mode that requires all components to pass, which can filter spurious results in the case of combinatorial barcoding. Candidates not resolved by the shortlist proceed to mutation-based neighborhood search or exhaustive matching.

RAD tracks per-barcode usage statistics across both the seed-indexed shortlist and the full broader whitelist, accepting a barcode as a correction target only if it has been independently observed in raw reads. Barcodes with counts dominated by corrections rather than direct observations are flagged and rejected (by default: corrections exceeding 80% of total observations once at least 10 total and 2 raw hits have accumulated), preventing a rare but valid barcode, such as one representing a low-abundance cell or an early capture event, from being silently reassigned to a more abundant neighbor.

### Whitelist Generation Algorithm

To identify barcodes from noisy long-read sequencing data, RAD first extracts putative barcodes by aligning a user-provided adapter and extracting the downstream *n* bases (default: 16). Next, RAD filters all extracted putative barcodes against a user-provided complete chemistry whitelist (for example, the full CellRanger barcode inclusion list). RAD first computes the normalized counts per million (ncpm) for each barcode as 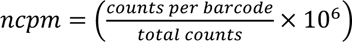, tabulates the log-normalized counts as ln(1 + *ncpm*), and then evaluates barcodes with log-normalized count greater than 2 (the “floor”).

The retained barcodes might still contain false positives. To filter them, RAD utilizes a Gaussian kernel-density estimate (KDE) over all retained barcodes’ count to find the threshold for identifying the true barcodes. If the KDE curve finds two or more peaks exist, RAD chooses the minimum (“saddle”) between the two tallest peaks. The saddle is extended to the leftmost plateau point (walking along the “flat valley”) to maximize the number of returned barcodes.

When a stable valley is absent, a positive-count Poisson distribution supplies a fallback threshold at its 95th percentile. The rate is set to the mean positive barcode count. A single-mode safeguard checks the position of this threshold relative to the peak and bandwidth. Separate density estimates for provisional high- and low-count populations provide a 50% mixture boundary. This boundary is diagnostic when a stable saddle is present and can refine the no-saddle fallback.

The initial “floor” is also revisited using an adjusted upper-tail hazard, calculated from the density and survival function of unique-barcode scores. A proposed lower boundary must be stable at 0.8, 1, and 1.25 times the density-estimation bandwidth. The spread between these boundaries must not exceed half a bandwidth. The candidate must have at least 5% local prominence and sufficient barcode support. Lowering the floor must add at least 50 barcodes and at least 1% of the original above-floor population, while retaining the required ordering of the hazard boundary, lower peak, saddle, and upper peak. Rescued barcodes must also satisfy the positive-count Poisson criterion.

### Analysis

#### Computational benchmarking

Computational speed and efficiency was benchmarked with the complete LongBench^11^ single nucleus ONT and single cell PB inputs, respectively. RAD, BLAZE, and Flexiplex were run with 4, 16, and 64 threads respectively. Each condition was repeated three times on nodes with AMD EPYC 7543 processors. Whitelist generation and demultiplexing were timed separately. Each demultiplexing run used its corresponding discovered whitelist. GNU time measured elapsed time and maximum resident set size for each phase. Flexiplex discovery included both discovery and filtering; their times were added, and the larger memory peak was retained. Total processing time was the sum of discovery and demultiplexing times within each repetition. Peak process memory was the GNU-time maximum resident set size across those phases.

#### Visium HD barcode recovery and spatial read counts

The long-read Visium HD mouse-brain dataset^18^ was compared with the matched Space Ranger spatial output. RAD extracted the UMI and the two barcode components according to the supplied read layout. Complete spatial assignments required both components. Barcode components were linked to spatial coordinates, and 2-µm coordinates were aggregated into 16-µm bins. Each binned coordinate was obtained by integer division of its original axis coordinate by eight. Space Ranger *in_tissue* annotations defined the displayed tissue reference. Barcode-associated counts were linked to the same spatial bins for visualization.

#### Synthetic data generation

Synthetic long-read sequences were obtained from the published ScTagger simulation^21^. The ScTagger-L simulation dataset contains approximately 5 million reads corresponding to 5,000 source barcodes. First, reads were shuffled to randomize their order within the file, and Q-scores were added. Base qualities were then sampled from the Badread^22^ Nanopore2023 quality model (random seed 20260731). For each read, all base quality scores were first sampled from the model’s overall distribution. The observed barcode was then aligned to its known source sequence to identify matching bases and sequence errors. New barcode quality scores were sampled from the Badread model distributions for these alignment patterns. These scores replaced the initial scores within the barcode region. If the barcode could not be located, the initial scores were retained for the entire read.

#### Comparing RAD, BLAZE, and Flexiplex performance on synthetic data

Whitelist generation was assessed by exact comparison of each selected set with the source barcode set. Both uncorrected raw barcodes and final barcodes were collected from the output of each tool. Briefly, RAD tagged each read with the final barcode as CB (cell identifier), and the uncorrected raw sequence as CR (uncorrected barcode sequence bases), directly in the output header using standard SAM formatting^45^. BLAZE stores its raw extractions in an auxiliary file (*putative_bc.csv*), which was paired against the final header-encoded barcode. For both BLAZE and RAD, read headers were extracted and parsed using *seqkit*^46^. As Flexiplex does not output the uncorrected raw output as a default, a custom build of Flexiplex (v1.02.6) was utilized. This build additionally returned a diagnostic assignment table that stored each read’s original identified, extracted barcode before correction, final barcode, barcode/flank edit distances, and the UMI. A separate Python script was utilized to tabulate the results. Barcodes were classified as “corrected” when the raw and final barcodes differed, and as “incorrect” when the final barcode did not match the ScTagger ground truth. Sensitivity – the fraction of detected true over total barcodes, precision – the fraction of true barcodes in the generated whitelist, and F1 score – the harmonic mean of sensitivity and precision, were evaluated for each tool.

#### Benchmarking off-list read annotations

Off-list correction, or the propensity to overcorrect barcodes during demultiplexing, was measured by supplying each tool with an incorrect, non-overlapping short-read size-matched whitelist. ScTagger-L and LongBench data originated from the 10x 3’ v3 inclusion list, while RAGE-seq data originated from the 10x 3’ v2 inclusion list. While ScTagger-L had a known ground-truth due to its synthetic nature (n = 5,000 total barcodes), the filtered short-read CellRanger cell barcodes were utilized as ground truth for both RAGE-seq (n = 4,505) and LongBench (n = 5,211). For all three experiments, the known ground truth barcodes were removed from the global inclusion lists. Target off-lists were then sampled independently without replacement using the Python 3.9.16 random.Random command, from the remainder of the whitelist. This process was repeated three times to generate technical replicates. Within each replicate, all three tools received the same input reads and off-list. For this experiment, LongBench was subsampled to 10 million reads with Seqkit 2.8.2. For assignment, all tools were allowed a maximum barcode edit distance of 2; Flexiplex was allowed a flank edit distance of up to 8.

#### Assessing failure modes in demultiplexing in the LongBench dataset

Barcode correction was evaluated across the LongBench single-cell and single-nucleus ONT and PacBio datasets to measure how correction redistributed reads among cell barcode identities. The complete chemistry whitelist (*3M-february-2018-3v3.txt*) defined all valid barcode sequences. Matched CellRanger-filtered barcodes defined the reference cell population. Barcodes were classified as “corrected” when the raw, extracted barcode sequence differed from the final assignment. Raw barcodes not originating from the complete chemistry whitelist and corrected to barcodes within the reference cell population, were termed as *corrected off-list barcodes*. These were considered as properly corrected. We then examined raw barcodes that did originate from the complete whitelist but were improperly corrected to a barcode within the reference cell population. If the improperly corrected raw barcode originated from the reference cell population, it was counted under failure mode 1. If the raw barcode originated from the complete chemistry whitelist, it was counted under failure mode 2. For assignment, all tools were allowed a maximum barcode edit distance of 2. Flexiplex utilized a flank edit distance of 0 for whitelist generation and 8 during assignment. For single-cell datasets, BLAZE was run with an – expect-cells parameter of 5,000. As previously reported for the single-nuclei datasets^47^, BLAZE was run with a –force-cells parameter of 10,000.

#### Whitelist generation benchmarking

The 37 datasets used (see Data Availability) and exact commands (see Code Availability) used to generate whitelists are provided. Summary statistics were generated with seqkit stats^46^. Whitelist generation for all datasets used RAD with default settings. BLAZE was provided the approximate number of short-read whitelist barcodes in the dataset for its –expect-cells parameter. RAD 1.0.2 was run on each complete input with a maximum adapter error of 0.3; each dataset used its corresponding adapter sequence. The LongBench single-nucleus ONT and PB datasets used –force-cells 10,000. Flexiplex was used to generate the barcode whitelist with flanking adapter distance (flag –f) set to either 0 or 8, followed by flexiplex-filter; the best result of the two runs was used.

#### Determination of immune cell type from RAGE-seq data

To assign receptor identity prior to demultiplexing, raw reads were aligned directly to the TRUST4^48^ BCR/TCR reference file, which contains curated immunoglobulin (IG) and T-cell receptor (TCR) gene segments from the human genome (GRch38^49^). Reads were aligned with *minimap2*^10^ using the default long-read mapping parameters for Nanopore sequencing (-ax *map-ont*). Receptor identity was determined according to the reference sequence of the best alignment; reads mapping to IG segments were labeled as BCR, reads mapping to TR segments were labeled as TCR, and all others were classified as unmapped. The resulting data was stored as a .csv file containing the original read ID and read identity.

#### Benchmarking FLAMESv2 against RAD on data from the LR-Split-seq protocol

RAD and FLAMESv2 2.7.1 processed reads from the same C2C12 LR-Split-seq dataset^8^ utilized in the original FLAMESv2 pre-print for benchmarking^35^. FLAMESv2 was run following its parameters from the original work^50^. RAD was run with its default settings for joint barcodes, with a maximum correction edit distance of 2. Both tools received the complete barcode catalog for each of the three independent barcodes. After demultiplexing, the complete set of barcodes were used to match assignments to the original 568 short-read cell identities^8^. UMI sequences were counted per cell and compared against the published short-read UMI counts across all three cell lines.

#### Genotype-based cell-line assignment in LongBench

Following the protocol from the original work, a fixed reference genotype panel was constructed from separate bulk Illumina RNA-sequencing libraries for the eight cancer cell lines. Bulk reads were processed with *fastp* 0.23.4^51^ and uniquely aligned to the GENCODE v44 human primary assembly using annotation-assisted Rsubread *subjunc* mapping^52^. CellSNP-lite 1.2.3^53^ called genotypes at candidate sites from the 1000 Genomes phase-3 hg38 panel with population allele frequency greater than 5%. The resulting reference panel was validated by re-analyzing the released Illumina single-nucleus allele-count matrices with Vireo 0.5.8^54^ and comparing the resulting classifications with the published assignments.

The complete ONT single-nuclei read sets demultiplexed by RAD, BLAZE, and Flexiplex were aligned using the FLAMES 2.0.1^35^ genome-alignment workflow with *minimap2*. Corrected cell barcodes and UMIs were retained in the corresponding SAM-formatted^45^ alignment tags (CB and UB, respectively). CellSNP-lite counted alleles using each tool’s selected barcode list and the common reference panel, with a minimum minor allele frequency of 0.1, a minimum total site count of 10, and genotype calling enabled. Vireo used the reference genotype calls with ambient-RNA estimation to classify each nucleus as a named cell-line singlet, doublet, or un-annotated.

Long-read classifications were then matched to the released Illumina reference by barcode sequence. Each Illumina reference singlet was classified as assigned to the same cell line, assigned to a different cell line, doublet, un-annotated, or not detected. Not detected denoted a reference barcode absent from the tool’s selected whitelist. Cell-line identity recall was calculated as the proportion of reference singlets assigned to the same cell line, both overall and separately for each cell line. Conditional concordance was calculated as the number of same-line assignments divided by the combined number of same-line and different-line assignments.

Genotype classifications were displayed on a common UMAP constructed from the released Illumina single-nucleus count matrix. All barcodes in the filtered matrix were retained. Counts were normalized to 10,000 per nucleus and log-transformed using Seurat^55^ LogNormalize. The 3,000 variable genes selected with the *vst* method were scaled and used to calculate 50 principal components. UMAP used the first 30 principal components, 30 neighbors, a minimum distance of 0.3, cosine distance, and random seed 20260805. Each tool’s classifications were overlaid on the same barcode-matched coordinates.

#### Differential Cell Line Expression Analysis in LongBench Single-Nuclei Data

Differential expression was assessed using the LongBench single nucleus ONT dataset. Each analysis included nuclei assigned by Vireo to the target, or comparator, cell-line group. Doublets and unassigned nuclei were excluded.

We utilized the paired short-read Illumina gene expression data as a reference and restricted analysis to the 8,135 genes eligible across all four processing pipelines. Eligibility was defined by a minimum of 10 reads per gene, and detection in at least 1% of queried cells. Counts were normalized as 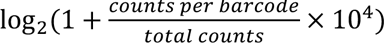. One-versus-rest differential-expression calls required an adjusted P value below 0.05, detection in at least 10% of nuclei in either group, and an absolute log₂ fold change of at least 0.5.

For comparisons benchmarked against Illumina, eligible genes were intersected across the Illumina, RAD, BLAZE, and Flexiplex workflows. Illumina reference calls required an adjusted P-value below 0.05 and an absolute Δ log_!_ *FC* ≥ 1. Genes enriched in each comparison group were evaluated separately. Precision was the proportion of long-read differential-expression calls that matched the Illumina reference in direction, sensitivity, or recall, was the proportion of same-direction Illumina reference calls recovered by the long-read workflow, and F1 was the harmonic mean of precision and recall.

## Data Availability

The ScTagger-S and -L source simulations are available from Figshare at <u>10.6084/m9.figshare.19740475.v1</u>. ScMixology2 (**<u>ERR9958136</u>**) and iPSC-derived neural progenitor cells (PromethION, **<u>ERR9958135</u>**) used in the BLAZE benchmarks are available from ENA (European Nucleotide Archive) under **<u>PRJEB54718</u>**. Published barcode tables and matched CellRanger reference data for these benchmarks are available from Figshare at doi:10.26188/20335203.v1. LR-Split-seq long-read data are available at **<u>SRR13948564</u>**, while corresponding short-read data and cell identities are available at **<u>SRR13948565</u>**. LongBench single-cell and single-nucleus ONT and PacBio data, matched Illumina data, and bulk Illumina data from the eight constituent cell lines are available under **<u>GSE303762</u>** and **<u>PRJNA1171488</u>**. RAGE-seq cell-line mixture and tumor-draining lymph-node data (TDLN) are available under ENA accession **<u>PRJEB28878</u>**. The P7, P8 VDJ-enriched, and P8 whole-transcriptome datasets from Beaulaurier *et al.*^29^ are available under **<u>PRJNA1087553</u>**. Human hippocampal long-read data from Joglekar *et al.*^28^ are available through NeMO collection <u>dat-717krsa</u>, within the <u>human single-nucleus data directories</u>. The Isosceles ovarian cancer cell-line mixture and matched Illumina data are available under **<u>GSE248115</u>.** MPNST long-read data are available under **<u>PRJEB90127</u>** and **<u>PRJEB77277</u>**. Mouse olfactory bulb spatial transcriptomic data are available under **<u>GSE153859</u>.** The four-flow-cell Visium HD 3′ mouse-brain dataset data are available through the <u>Oxford Nanopore Open Data release, visium_hd_2025.06</u>. Human tonsil sequencing data from Engblom *et al.*^32^ are available through EGA dataset **<u>EGAD00001011062</u>** (study accession **<u>EGAS00001007337</u>**). Access requires approval from the Karolinska Institutet Data Access Committee (**<u>EGAC00001003294</u>**) under the dataset’s access agreement and ethics requirements. Associated processed data, metadata, tissue images, and analysis scripts are publicly available at doi:10.5281/zenodo.7961605.Comparative benchmarking across all three tools presented in **Figure 4** included each of these datasets, excepting Visium HD and LR-Split-seq.

## Code Availability

Source code for RAD, along with all read layout .csv files and downstream analysis scripts required to reproduce the analyses in this manuscript, is available at https://github.com/indianewok/rad. Running commands and analysis code for this study are available at https://github.com/indianewok/rad-manuscript.

## AI Disclosure

Both Claude and Codex were utilized to assist with codebase maintenance and end-to-end Github documentation, repository set-up and clean-up, data cleaning and deduplication, consolidation of methods, early drafting of the portions of the manuscript, and orchestrating workflows for large benchmarking tasks. The authors manually reviewed, revised, and assessed results generated from all AI-assisted code for consistency and correctness. The authors take full responsibility for the final work.

## Acknowledgements

The authors would like to thank Dr. Aaron McKenna and Dr. Kenneth Hoehn for helpful discussions and for providing computational resources throughout the development of this project. The authors would also like to thank Dr. Hunter Melton and Dr. Celeste Parra Bravo for their thoughtful conversations in developing this tool. Figure 3D created in BioRender. Ackerman, M. (2026) https://BioRender.com/ic2r73i

## Author contributions

C.M.V., L.S., and M.E.A. conceived the project. C.M.V. developed the pipeline and performed the analyses. C.M.V., M.C., Y.H., L.S., and M.E.A. analyzed the data. C.M.V. and M.C. wrote the initial manuscript draft. All authors evaluated results and contributed to the manuscript editing.

## References

1. Marx, V. Method of the year: long-read sequencing. Nat Methods 20, 6–11 (2023).

2. Dijk, E. L. van, Jaszczyszyn, Y., Naquin, D. & Thermes, C. The Third Revolution in Sequencing Technology. Trends in Genetics 34, 666–681 (2018).

3. Marx, V. Long road to long-read assembly. Nat Methods 18, 125–129 (2021).

4. Tian, L. et al. Comprehensive characterization of single-cell full-length isoforms in human and mouse with long-read sequencing. Genome Biol 22, 310 (2021).

5. Kovaka, S., Ou, S., Jenike, K. M. & Schatz, M. C. Approaching complete genomes, transcriptomes and epi-omes with accurate long-read sequencing. Nat Methods 20, 12–16 (2023).

6. Jain, C., Rhie, A., Hansen, N. F., Koren, S. & Phillippy, A. M. Long-read mapping to repetitive reference sequences using Winnowmap2. Nat Methods 19, 705–710 (2022).

7. Vaz-Drago, R., Custódio, N. & Carmo-Fonseca, M. Deep intronic mutations and human disease. Hum Genet 136, 1093–1111 (2017).

8. Rebboah, E. et al. Mapping and modeling the genomic basis of differential RNA isoform expression at single-cell resolution with LR-Split-seq. Genome Biology 22, 286 (2021).

9. Veiga, D. F. T. et al. A comprehensive long-read isoform analysis platform and sequencing resource for breast cancer. Science Advances 8, eabg6711 (2022).

10. Li, H. Minimap2: pairwise alignment for nucleotide sequences. Bioinformatics 34, 3094–3100 (2018).

11. You, Y., et al. Benchmarking long-read RNA-sequencing technologies with LongBench: a cross-platform reference dataset profiling cancer cell lines with bulk and single-cell approaches. Preprint at 10.1101/2025.09.11.675724 (2026).

12. Gupta, P., O’Neill, H., Wolvetang, E. J., Chatterjee, A. & Gupta, I. Advances in single-cell long-read sequencing technologies. NAR Genom Bioinform 6, lqae047 (2024).

13. Fu, Y. et al. Single cell and spatial alternative splicing analysis with Nanopore long read sequencing. Nat Commun 16, 6654 (2025).

14. Al’Khafaji, A. M. et al. High-throughput RNA isoform sequencing using programmed cDNA concatenation. Nat. Biotechnol. 42, 582–586 (2024).

15. Cheng, O. et al. Flexiplex: a versatile demultiplexer and search tool for omics data. Bioinformatics 40, btae102 (2024).

16. Kuijpers, L. et al. Split Pool Ligation-based Single-cell Transcriptome sequencing (SPLiT-seq) data processing pipeline comparison. BMC Genomics 25, 361 (2024).

17. You, Y. et al. Identification of cell barcodes from long-read single-cell RNA-seq with BLAZE. Genome Biology 24, 66 (2023).

18. Visium HD 3’ mouse brain. EPI2ME https://epi2me.nanoporetech.com/visium_hd_2025.06/ (2025).

19. epi2me-labs/wf-single-cell. EPI2ME (2026).

20. Martin, M. Cutadapt removes adapter sequences from high-throughput sequencing reads. EMBnet.journal 17, 10–12 (2011).

21. Ebrahimi, G. et al. Fast and accurate matching of cellular barcodes across short-reads and long-reads of single-cell RNA-seq experiments. iScience 25, 104530 (2022).

22. Wick, R. Badread: simulation of error-prone long reads. JOSS 4, 1316 (2019).

23. Cell Ranger’s Gene Expression Algorithm | Official 10x Genomics Support. 10x Genomics https://www.10xgenomics.com/support/software/cell-ranger/10.0/algorithms-overview/cr-gex-algorithm.

24. Riemondy, K. A. et al. Recovery and analysis of transcriptome subsets from pooled single-cell RNA-seq libraries. Nucleic Acids Research 47, e20–e20 (2019).

25. Zhao, L., Liu, Z., Levy, S. F. & Wu, S. Bartender: a fast and accurate clustering algorithm to count barcode reads. Bioinformatics 34, 739–747 (2018).

26. Petukhov, V. et al. dropEst: pipeline for accurate estimation of molecular counts in droplet-based single-cell RNA-seq experiments. Genome Biol 19, 78 (2018).

27. Singh, M. et al. High-throughput targeted long-read single cell sequencing reveals the clonal and transcriptional landscape of lymphocytes. Nat Commun 10, 3120 (2019).

28. Joglekar, A. et al. Single-cell long-read sequencing-based mapping reveals specialized splicing patterns in developing and adult mouse and human brain. Nature Neuroscience 27, 1051–1063 (2024).

29. Beaulaurier, J. et al. De novo antibody identification in human blood from full-length single B cell transcriptomics and matching haplotype-resolved germline assemblies. Genome Res 35, 929–941 (2025).

30. Hamraoui, A., et al. A systematic benchmark of bioinformatics methods for single-cell and spatial RNA-seq nanopore long reads data. NAR Genom. Bioinform. 8, lqag070 (2026).

31. Kabza, M. et al. Accurate long-read transcript discovery and quantification at single-cell, pseudo-bulk and bulk resolution with Isosceles. Nature Communications 15, 7316 (2024).

32. Engblom, C. et al. Spatial transcriptomics of B cell and T cell receptors reveals lymphocyte clonal dynamics. Science 382, eadf8486 (2023).

33. Lebrigand, K. et al. The spatial landscape of gene expression isoforms in tissue sections. Nucleic Acids Research 51, e47 (2023).

34. Long-read transcriptomics - opportunities and challenges | npj Genomic Medicine. https://www.nature.com/articles/s41525-026-00617-5.

35. Wang, C. et al. Igniting full-length isoform analysis in single-cell and spatial RNA-seq data with FLAMESv2. 2025.10.19.683327 Preprint at 10.1101/2025.10.19.683327 (2026).

36. Huang, Y.-H. et al. POU2F3 is a master regulator of a tuft cell-like variant of small cell lung cancer. Genes Dev. 32, 915–928 (2018).

37. Rudin, C. M. et al. Molecular subtypes of small cell lung cancer: a synthesis of human and mouse model data. Nat. Rev. Cancer 19, 289–297 (2019).

38. Wooten, D. J. et al. Systems-level network modeling of Small Cell Lung Cancer subtypes identifies master regulators and destabilizers. PLoS Comput. Biol. 15, e1007343 (2019).

39. Calis, J. J. A. & Rosenberg, B. R. Characterizing immune repertoires by high throughput sequencing: strategies and applications. Trends in Immunology 35, 581–590 (2014).

40. Shlesinger, D. et al. Single-cell immune repertoire sequencing of B and T cells in murine models of infection and autoimmunity. Genes Immun 23, 183–195 (2022).

41. Stavnezer, J., Guikema, J. E. J. & Schrader, C. E. Mechanism and Regulation of Class Switch Recombination. Annual Review of Immunology 26, 261–292 (2008).

42. Šošić, M. & Šikić, M. Edlib: a C/C ++ library for fast, exact sequence alignment using edit distance. Bioinformatics 33, 1394–1395 (2017).

43. Zhao, M., Lee, W.-P., Garrison, E. P. & Marth, G. T. SSW Library: An SIMD Smith-Waterman C/C++ Library for Use in Genomic Applications. PLoS ONE 8, e82138 (2013).

44. Myers, G. A Fast Bit-Vector Algorithm for Approximate String Matching Based on Dynamic Programming. Journal of the ACM 46, 395–415 (1999).

45. samtools/hts-specs: Specifications of SAM/BAM and related high-throughput sequencing file formats. https://github.com/samtools/hts-specs.

46. Shen, W., Le, S., Li, Y. & Hu, F. SeqKit: A Cross-Platform and Ultrafast Toolkit for FASTA/Q File Manipulation. PLoS ONE 11, e0163962 (2016).

47. LongBench_workflow/config/config_lr_sc_sn.yaml at 37a9506df7bcd92c488c0e4b2a5cbe09fa1fc19d · youyupei/LongBench_workflow. https://github.com/youyupei/LongBench_workflow/blob/37a9506df7bcd92c488c0e4b2a5cbe09fa1fc19d/config/config_lr_sc_sn.yaml#L45-L73.

48. Song, L. et al. TRUST4: immune repertoire reconstruction from bulk and single-cell RNA-seq data. Nat Methods 18, 627–630 (2021).

49. Homo sapiens genome assembly GRCh38. NCBI https://www.ncbi.nlm.nih.gov/datasets/genome/GCF_000001405.26/.

50. rad-manuscript/source_data/figure_05/provenance/flames_paper_snapshot/upstream/ChangqingW_FLAMES_figures/files/config_splitseq.json at flames-configs-2026-10-30 · indianewok/rad-manuscript · GitHub. https://github.com/indianewok/rad-manuscript/blob/flames-configs-2026-10-30/source_data/figure_05/provenance/flames_paper_snapshot/upstream/ChangqingW_FLAMES_figures/files/config_splitseq.json#L21-L71.

51. Chen, S., Zhou, Y., Chen, Y. & Gu, J. fastp: an ultra-fast all-in-one FASTQ preprocessor. Bioinformatics 34, i884–i890 (2018).

52. Liao, Y., Smyth, G. K. & Shi, W. The R package Rsubread is easier, faster, cheaper and better for alignment and quantification of RNA sequencing reads. Nucleic Acids Research 47, e47 (2019).

53. Huang, X. & Huang, Y. Cellsnp-lite: an efficient tool for genotyping single cells. Bioinformatics 37, 4569–4571 (2021).

54. Huang, Y., McCarthy, D. J. & Stegle, O. Vireo: Bayesian demultiplexing of pooled single-cell RNA-seq data without genotype reference. Genome Biology 20, 273 (2019).

55. Hao, Y. et al. Dictionary learning for integrative, multimodal and scalable single-cell analysis. Nat Biotechnol 42, 293–304 (2024).

